# Heritability estimates of antler and body traits in white-tailed deer (*Odocoileus virginianus*) from genomic-relatedness matrices

**DOI:** 10.1101/2020.04.22.053165

**Authors:** Aidan Jamieson, Spencer J Anderson, Jérémie Fuller, Steeve D Côté, Joseph M Northrup, Aaron BA Shafer

## Abstract

Estimating heritability (*h*^2^) is required to predict the response to selection and is useful in species that are managed or farmed using trait information. Estimating *h*^2^ in free-ranging populations is challenging due to the need for pedigrees; genomic-relatedness matrices (GRMs) circumvent this need and can be implemented in nearly any system where phenotypic and SNP data are available. We estimated the heritability of five body and three antler traits in a free-ranging population of white-tailed deer (*Odocoileus virginianus*) on Anticosti Island, Quebec, Canada. We generated GRMs from >10,000 SNPs: dressed body mass and peroneus muscle mass had moderate *h*^2^ values of 0.49 and 0.56, respectively. Heritability in male-only antler features ranged from 0.00 to 0.51 and had high standard errors. We explored the influence of minor allele frequency and data completion filters on *h*^2^: GRMs derived from fewer SNPs had reduced *h*^2^ estimates and the relatedness coefficients significantly deviated from those generated with more SNPs. As a corollary, we discussed limitations to the application of GRMs in the wild, notably how skewed GRMs increase variance around *h*^2^ estimates. This is the first study to estimate *h*^2^ on a free-ranging population of white-tailed deer and should be informative for breeding designs and management.

---

How individuals respond to selection is dependent, in part, on the heritability (*h*^2^) of the focal traits (Wright, 1920; Falconer and Mackay, 1996; Gillespie, 2004). Quantifying *h*^2^ is essential to predicting adaptive responses (e.g. Dziedzic *et al*., 2019), opening the potential for gene targeted management and conservation (Kardos and Shafer, 2018). Classically, *h*^2^ was estimated by comparing the phenotypic variance (*V*_P_) of known relatives, which allowed for estimating additive genetic variance (*V*_A_) with narrow-sense heritability being *h*^2^ = *V*_A_ / *V*_P_ and Visscher, 2008). In free-ranging populations this estimation has required the construction of pedigrees that were then integrated into a mixed-effects model, known as the animal model (Wilson *et al*., 2010). The animal model disentangles *V*_P_ into its additive (*V*_A_) and environmental components, enabling *h* to be estimated.

The tracking of individuals and construction of pedigrees in free-ranging populations is time-consuming and costly (Clutton Brock and Sheldon, 2010). As a result, alternative pedigree-free approaches have emerged in which molecular genealogies can be structured into relatedness matrices that replace the pedigree in an animal model (Frentiu *et al*., 2008). These genealogies have relied on microsatellites, which can be problematic due to biased estimates in small sample sizes (Wang, 2017) and the recommended number of loci exceeding that used in most studies (Kopps *et al*., 2015). This changed with the advent of high-throughput sequencing that made the generation of thousands of single nucleotide polymorphisms (SNPs) feasible for most organisms (Andrews *et al*., 2016). Genomic-relatedness matrices (GRM) can be generated and used to estimate the total amount of *V*_P_ captured by the panel of SNPs (Yang *et al*., 2011; Stanton-Geddes *et al*., 2013). Here, *V*_A_ is estimated across SNPs (see Ritland, 1996) and has produced *h*^2^ estimates similar to those of sibling studies (Yang *et al*., 2010). Though few, recent applications to wild populations have paid special attention to the impact of model choice and influence of missing data and minor allele frequencies (MAF) on number of retained SNPs and *h*^2^ estimates (e.g. Perrier *et al*., 2018; Gervais *et al*., 2019).

Quantifying *h*^2^ of specific traits in wild species that are farmed or harvested can be valuable for breeding design (Guo *et al*., 2018) and management decisions (Coltman, 2008). White-tailed deer (*Odocoileus virginianus*) are a big game species with economic and cultural values across North America. Annual revenue from hunting is in the billions (Cambronne, 2013) and close to 50 million USD from deer farming (see https://www.nass.usda.gov/AgCensus/). Antler and body size are positively correlated with male reproductive success (Newbolt *et al*., 2017), and *h*^2^ for these traits have been estimated in captively bred populations. For example, *h*^2^ was estimated using parent-offspring trios at 0.32 for the number of antler points and for0.45 main beam length, though with high standard errors (Michel *et al*., 2016). Similarly, *h*^2^ was estimated between 0.58 and 0.64 for body mass (Williams *et al*., 1994). Recent work suggests a highly polygenic nature of these traits (Anderson *et al*., 2019), but *h*^2^ has yet to be quantified in a free-ranging population of white-tailed deer. Theoretical and empirical evidence suggests that bottlenecked populations, such as those we might expect to see in captivity, have increased *V*_A_ (Goodnight, 1988; Whitlock *et al*., 1993; van Heerwaarden *et al*., 2008; Jarvis *et al*., 2011; Taft and Roff, 2012), thus it is unclear how *h*^2^ values transfer from captivity to the wild. Here, we take advantage of an extensive phenotypic database and DNA archive to estimate *h*^2^ in antler and body traits of free-ranging white-tailed deer using GRMs.

## Materials and Methods

### Individual Sampling and phenotypes

Ear or muscle tissue was obtained and preserved in 95% ethanol between 2003 and 2014 from harvested white-tailed deer on Anticosti Island, Quebec, Canada (Figure S1). This population originated from ~220 individuals introduced in the late 1800s and shows no evidence of a founder effect (Fuller *et al*., 2020). All individuals had an extensive set of phenotypic metrics taken (Table S1), were sexed, and aged using cementum layers in incisor teeth (Hamlin *et al*., 2000) with only animals >1.5 years analyzed. Phenotypes were assessed for correlation and only one was retained if Spearman’s *r* was > 0.70. DNA was isolated from the tissue using a phenol-chlorophorm-isoamyl procedure and stored at −80 °C.

### Genome relatedness matrix and genotyping

We used the SNP data generated in Fuller *et al*., (2020) that applied a rigorous filtering approach to generate 13,420 SNPs. We took this VCF file and further filtered it to include only scaffolds that had >1 SNP present, and made a separate data file that only included males. To quantify the impact of loci missingness (LM) and MAF filters on downstream inferences (e.g. Shafer *et al*., 2017) we generated subsets of data with 70, 80, 90 and 100% completion as well 1, 5 and 10% MAF requirements. The SNP genotypes were converted into a binary fileset using PLINK v.19 (Purcell *et al*., 2007) that was used to generate a GRM in GCTA v1.92.4 (Yang *et al*., 2011). The GRMs were input to a mixed-effects linear model that included year of harvest, age, and sex with a restricted maximum-likelihood (REML) approach to estimate *h*^2^ (Meyer, 1989). An F test of equality of variances was conducted in R v. 3.6.3. to test whether the GRM distributions differed among the filtered datasets, and we tested whether deviations from a 1:1 ratio, calculated as abs(1-F statistic), correlated to differences in the number of SNPs used to build the GRMs.

For a subset of male samples we genotyped the SRP54 locus that was previously found to be associated with antler morphology (Anderson *et al*., 2019). We custom designed a rhAmp SNP assay and genotyped individuals via qPCR with a QuantStudio 3 Real-Time PCR System (Thermo Fisher Scientific, Waltham, MA, USA). The thermocycling condition was 60 °C for 30 seconds followed by 95 °C for 10 minutes, then 40 cycles of 95 °C for 10 seconds, 60 °C for 30 seconds, and 68 °C for 20 seconds, followed by a final 30 seconds at 60 °C. The qPCR had a 5 μL reaction volume in each 0.1 mL well of a sealed 96-well optical plate. As above the GRMs were fitted to include year of harvest, age, and the SRP54 locus in an additive framework (0, 1, or 2 alleles).

Scripts have been uploaded to a GIT repository: https://gitlab.com/WiDGeT_TrentU

## Results

We obtained samples from 219 male and 218 female deer. The number of SNPs retained for body size and antler estimates are provided in Table 1. Increased loci completeness requirements and higher MAF values reduced the number of SNPs used to build the GRM, with the final SNP datasets containing between 1,041 and 10,386 SNPs. The distribution of relatedness values in the GRM showed primarily unrelated individuals, though there was evidence of full-sibling or parent-offspring pairings (Figure 1). The filtered GRM distributions differed with increased MAF and LM filters that skewed the relatedness coefficients (Table 2, Figure 2). Variance between GRMs (i.e. abs(1-F-statistic)) and the difference in number of SNPs used to build the GRMs were highly correlated (*β* = 1.63e-05, *R*^2^ = 0.92, P < 0.001).

**Figure 1.**
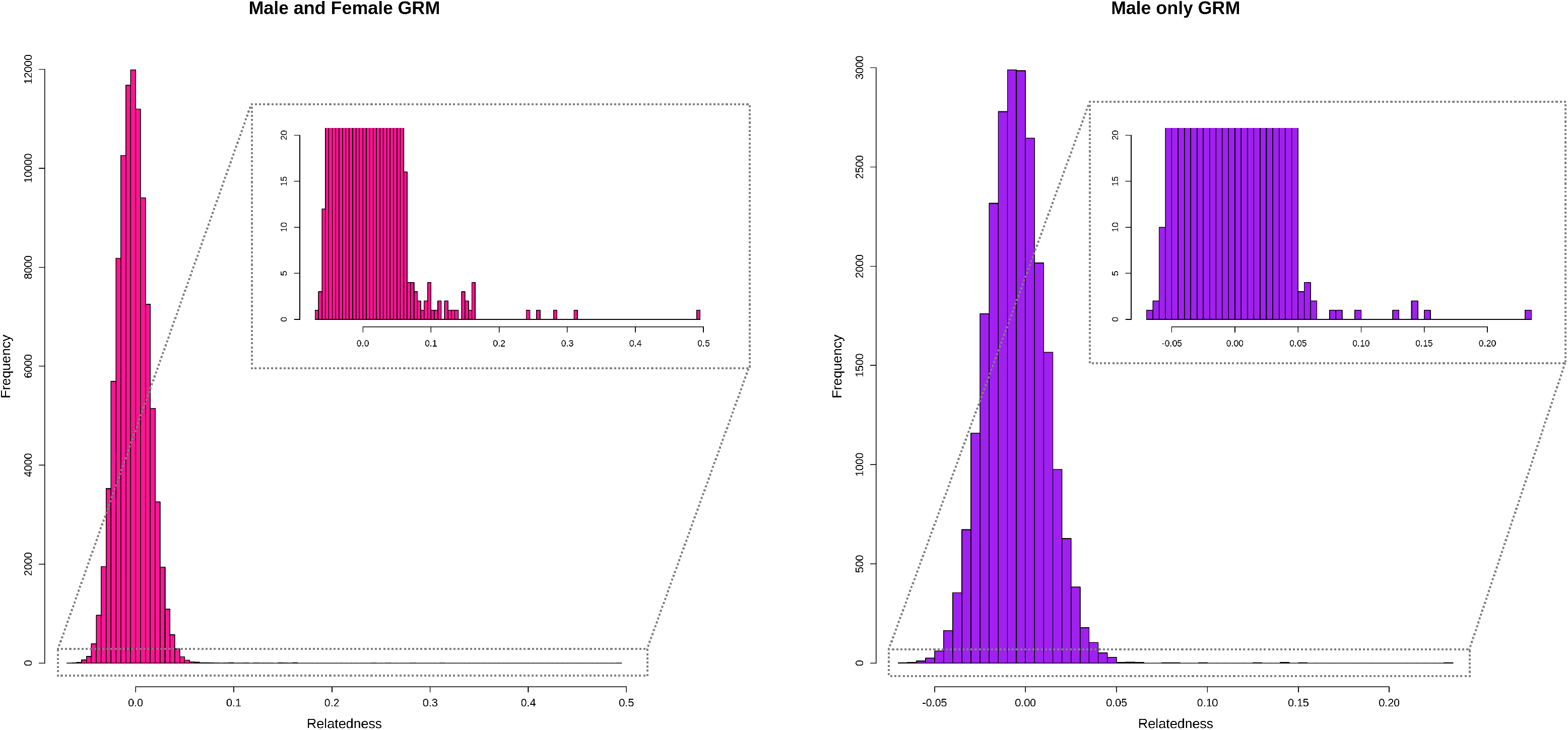
Genomic-relatedness matrices (GRMs) for white-tailed deer (*Odocoileus virginianus*) on Anticosti Island, Quebec, Canada. GRMs are divided into the entire dataset (male and female) and male only.

**Figure 2.**
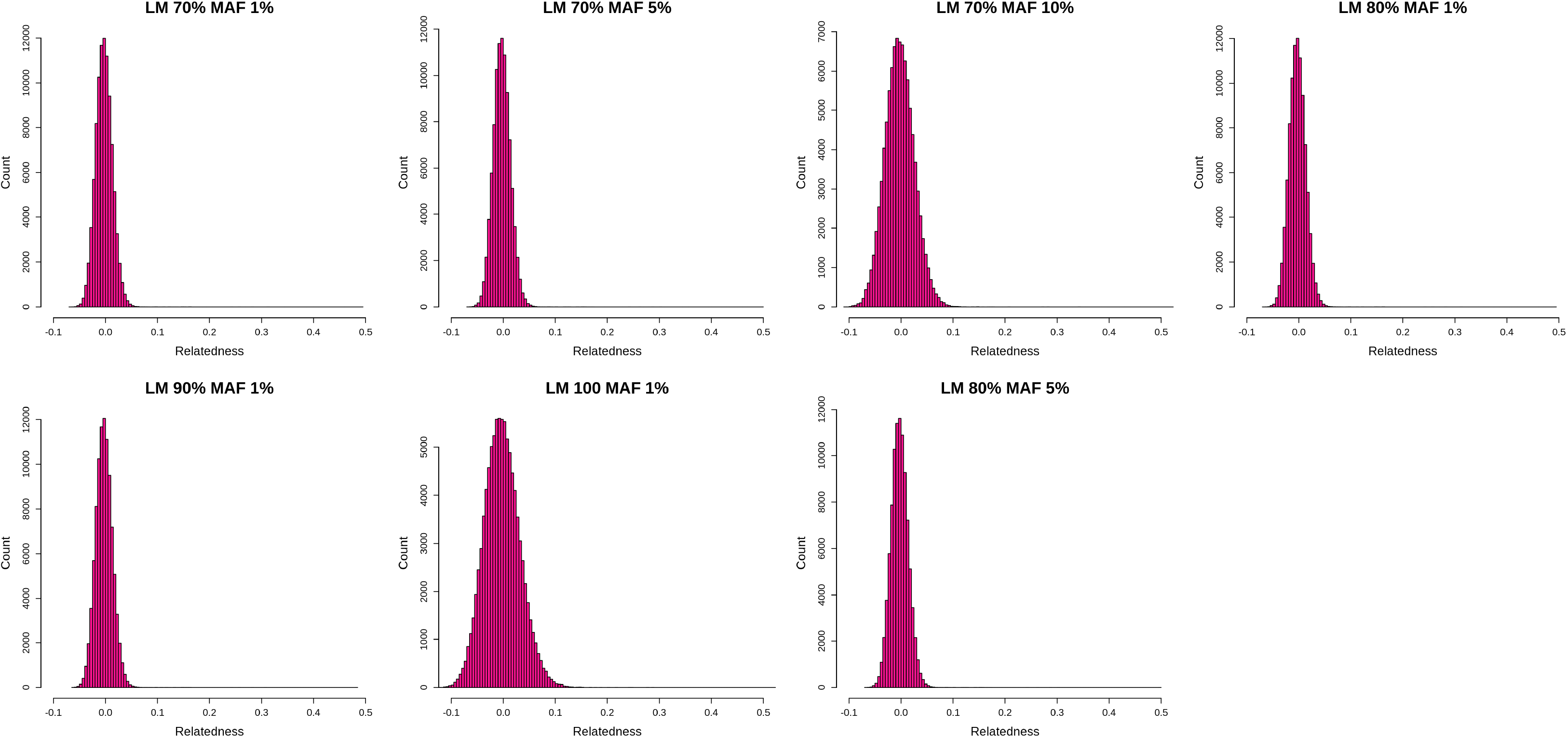
Genomic-relatedness matrices (GRMs) for the sampled white-tailed deer (*Odocoileus virginianus*). GRMs are from the entire dataset (male and female) and were generated with varying minor allele frequency (MAF) and loci missing (LM) filters.

**Table 1.**
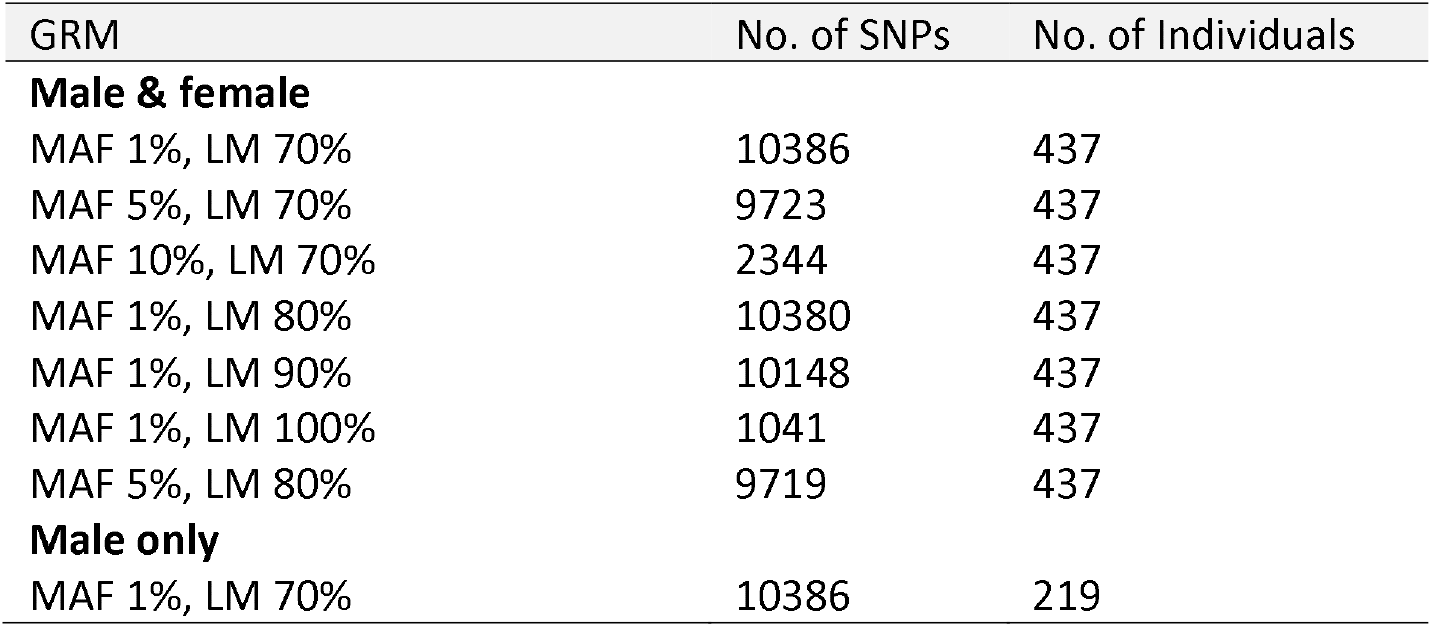
Number of SNPs and individuals retained in each genomic-relatedness matrix (GRM) after standard filtering. Additional filtering was only applied to full male & female datasets using different minor allele frequency (MAF) required loci completion (LM) criteria.

**Table 2.** Pairwise F tests of variances of the filtered genomic-relatedness matrices (GRM) in white-tailed deer (*Odocoileus virginanus*). We used different minor allele frequency (MAF) required loci completion (LM) criteria to generate GRMs. The F statistics and P values are presented, and *n* is 436. ΔSNPs refers to the difference in no. SNPs used to build the GRMs.

| GRM 1 | GRM 2 | ΔSNPs | F statistic | P value |
| --- | --- | --- | --- | --- |
| MAF 1%, LM 70% | MAF 5%, LM 70% | 663 | 0.99 | 0.38 |
| MAF 1%, LM 70% | MAF 10%, LM 70% | 8042 | 0.86 | <0.001 |
| MAF 1%, LM 70% | MAF 1%, LM 80% | 6 | 1.00 | 0.93 |
| MAF 1%, LM 70% | MAF 1%, LM 90% | 238 | 1.01 | 0.002 |
| MAF 1%, LM 70% | MAF 1%, LM 100% | 9345 | 0.88 | <0.001 |
| MAF 1%, LM 70% | MAF 5%, LM 80% | 667 | 1.00 | 0.41 |
| MAF 5%, LM 70% | MAF 10%, LM 70% | 7379 | 0.86 | <0.001 |
| MAF 5%, LM 70% | MAF 1%, LM 80% | 657 | 1.00 | 0.33 |
| MAF 5%, LM 70% | MAF 1%, LM 90% | 425 | 1.02 | <0.001 |
| MAF 5%, LM 70% | MAF 1%, LM 100% | 8682 | 0.88 | <0.001 |
| MAF 5%, LM 70% | MAF 5%, LM 80% | 4 | 1.00 | 0.95 |
| MAF 10%, LM 70% | MAF 1%, LM 80% | 8036 | 1.16 | <0.001 |
| MAF 10%, LM 70% | MAF 1%, LM 90% | 7804 | 1.17 | <0.001 |
| MAF 10%, LM 70% | MAF 1%, LM 100% | 1303 | 1.01 | 0.006 |
| MAF 10%, LM 70% | MAF 5%, LM 80% | 7375 | 1.15 | <0.001 |
| MAF 1%, LM 80% | MAF 1%, LM 90% | 232 | 1.01 | 0.004 |
| MAF 1%, LM 80% | MAF 1%, LM 100% | 9339 | 0.88 | <0.001 |
| MAF 1%, LM 80% | MAF 5%, LM 80% | 661 | 1.00 | 0.36 |
| MAF 1%, LM 90% | MAF 1%, LM 100% | 9107 | 0.86 | <0.001 |
| MAF 1%, LM 90% | MAF 5%, LM 80% | 429 | 0.98 | <0.001 |
| MAF 1%, LM 100% | MAF 5%, LM 80% | 8678 | 1.14 | <0.001 |

Dressed body mass and peroneus muscle mass for males and females together had heritability estimates of 0.49 and 0.56 respectively; all other body metrics had relatively large standard errors (SE; Table 3). Likewise, two male-only antler metrics had non-zero *h*^2^ estimates but high SE (Table 3). Smaller *h*^2^ estimates were observed from GRMs built with fewer SNPs (Table S2). Not surprisingly, age and sex had large effects on the body size metrics, with older individuals and males being larger on average; antler metrics also increased with age (Table 3). Estimates of *h*^2^ were lower when age and sex were not included in the model (Table S2), whereas removing year of harvest only had a minor effect (Tables S3; S4). We genotyped 74 males at the SRP54 locus (Dryad Accession XXXX); despite a positive effect size of this gene on antler metrics, all estimates had high SE precluding any meaningful inference (Table S5; S6).

**Table 3.**
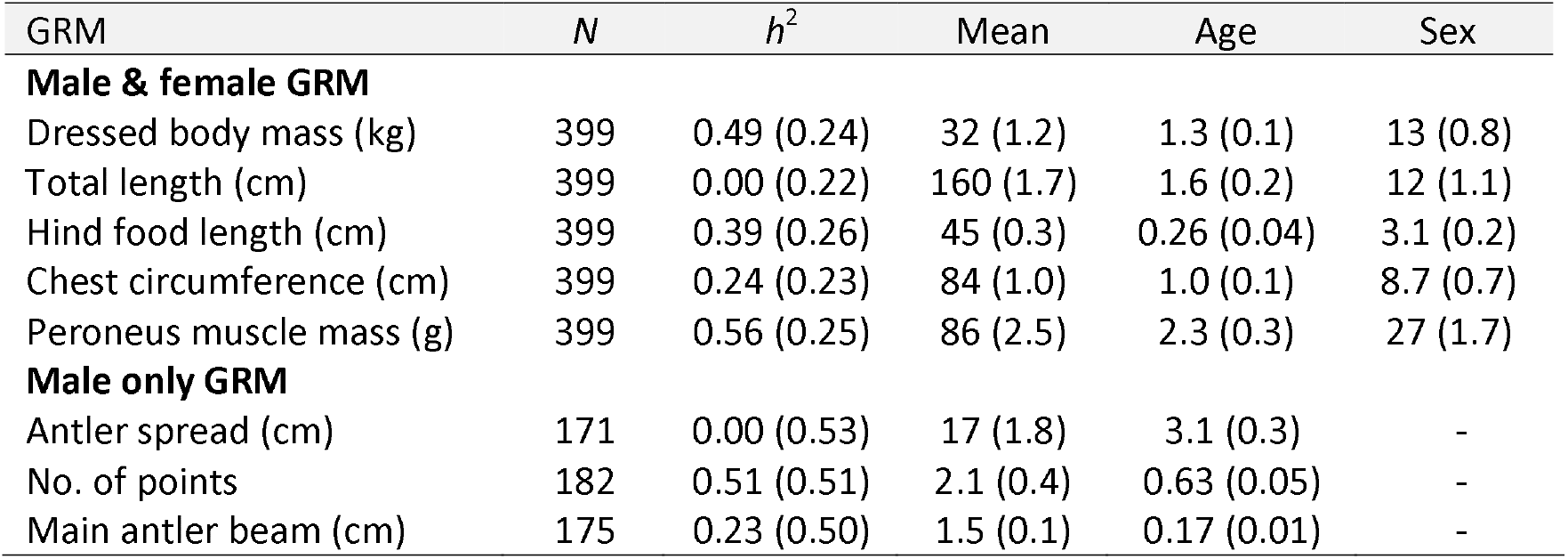
Estimation of heritability (*h*^2^) for body and antler traits in white-tailed deer (*Odocoileus virginianus*) on Anticosti Island, Quebec, Canada. The genomic-relatedness matrix (GRM) was generated using a minor allele frequency of 1% and required loci completion of 70%. Mean trait values and effect sizes of age and sex are provided but the effect of year is not shown. Standard errors for all metrics are included in parentheses. *N* refers to the number individuals retained after filtering for phenotype completeness.

## Discussion

White-tailed deer are an important big-game species that are hunted and farmed across North America. If larger antlered or bodied animals are preferentially harvested or bred, there is the potential for artificial selection provided such traits are heritable. Our data suggest that body and muscle mass are heritable (Table 2), which is consistent with most vertebrates (Mousseau and Roff, 1987) including deer (Williams *et al*., 1994). For antler metrics, despite *h*^2^ values similar to previous estimates (i.e. Lukefahr and Jacobson, 1998; Michel *et al*., 2016), our estimates had large standard errors; this is likely a reflection of the smaller male-only data set and a GRM skewed towards unrelated individuals (Figure 1) as captive studies suggest moderate levels of heritability. Whether harvest could induce artificial selection in white-tailed deer is an open question: some data have shown harvested antlers being larger than those naturally shed (Schoenebeck and Peterson, 2014; but see Ditchkoff *et al*., 2000), which could generate an evolutionary response. Still, non-genetic factors are large contributors to body size and antlers (Table 2) and in our view it is unlikely that harvest is a strong enough selective pressure to reduce antler size in most managed regions, given the generally large population size of deer, varied regulations, and often opportunistic nature of the hunt. Small and isolated populations, including farms, could see a response to selection, especially if *V*_A_ is higher (e.g. Taft and Roff, 2012).

The heritability of traits can vary by sex, age, and across seasons (Réale *et al*., 1999). This is an important consideration with GRM approaches for phenotypically stratified populations that despite circumventing the need for pedigrees, still require basic individual metadata. In our study this meant aging individuals through cross-sections of incisors (Hamlin et al. 2000), and not accounting for age and sex in the model resulted in downward biased *h*^2^ estimates (Table S2). Likewise, accounting for parturition date and litter size increased *h*^2^ estimates in captive white-tailed deer (Michel et al. 2016). Interestingly, including the year of harvest that presumably reflect environmental variation, particularly during summer, had a largely negligible effect on *h*^2^ (Table S3; S4). The relatively high standard errors surrounding *h*^2^, even after accounting for age and sex, is not uncommon as seen in Michel et al.’s (2016) analysis of captive deer, and more broadly ungulates (Thomas *et al*., 2002; Coltman *et al*., 2005). However, Sim and Coltman (2019) used GRMs and reported moderate *h*^2^ estimates with small SE of horn characteristics of thinhorn sheep (*Ovis dalli*): this difference is likely attributed to antlers being shed and regrown annually, and horn length being a more easily quantifiable metric.

Sample size, SNP number, and filtering criteria all must be considered when estimating *h*^2^ with GRMs (Visscher and Goddard, 2015; Gervais *et al*., 2019). Small sample size increases sampling variance, typically biasing *V*_A_ downwards (Villemereuil *et al*., 2013). With SNP data, the GRM variance is inversely proportional to that of *h*^2^ (Visscher and Goddard, 2015). Both our analysis and two other wild ungulate studies (Bérénos *et al*., 2014; Gervais *et al*., 2019) had GRMs skewed towards distant relatives; this undoubtedly played a role in *h*^2^ estimates, as sampling the full spectrum of relatives is difficult in wild populations. The use of SNP data also involves bioinformatics decisions. High MAF cutoffs should be avoided as this will shift the site-frequency spectrum towards more common alleles, resulting in reduced *h*^2^ estimates (Table S2; Gervais et al. 2019) and appreciably different GRMs (Table 2; Figure 2). Introducing a stringent loci missingness criteria in principle should improve estimates (Gervais et al. 2019), though we saw minimal evidence of this, likely due to the reduced SNP number (Table 1). Collectively, these filtering parameters all impact the quality and number of SNPs used to build the GRM, with wild vertebrate studies using a few thousand (Gervais *et al*., 2019; Sim and Coltman, 2019) to tens of thousands SNPs (Bérénos *et al*., 2014; Perrier *et al*., 2018). While the optimal sample size and SNP number will depend on demographic history and life history strategies of the focal population, the generation of GRMs and estimates of *h*^2^ in the wild opens up numerous basic and applied research opportunities, notably predicting the response to selection under harvest and climate change scenarios. In the case of white-tailed deer, we have shown that body size and likely antlers are moderately heritable and could respond to selection, which would be most likely in small and isolated populations.

## Acknowledgements

This work was supported by the Natural Sciences and Engineering Research Council of Canada Discovery Grants (ABAS and SDC); ComputeCanada Resources for Research Groups (ABAS); Canadian Foundation for Innovation: John R. Evans Leaders Fund (ABAS); and Industrial Chairs and Collaborative Research and Development Grants from the Natural Sciences and Engineering Research Council of Canada (SDC). We thank the outfitters of Anticosti Island and the Ministère des Forêts, de la Faune et des Parcs du Québec for logistical help associated with fieldwork, and all of the field assistants and technicians who collected samples and made phenotypic measurements. Special thanks to Louis Bernatchez for supporting the lab work.

