## Supplemental Tables and Figures for "Heritability estimates of antler and body traits in white-tailed deer (*Odocoileus virginianus*) from genomic-relatedness matrices"

**Figure S1.** Map of Anticosti Island, Quebec, Canada where white-tailed deer (*Odocoileus virginianus*) were sampled for this study.

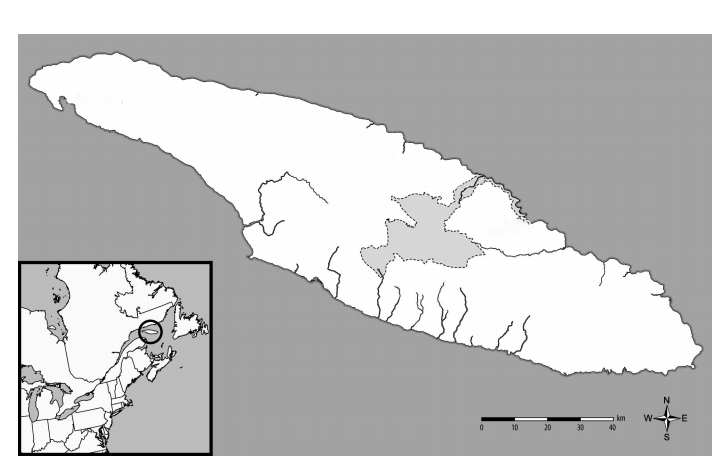

**Table S1.** Phenotypes of white-tailed deer measured in this study

Dressed body mass (kg)

Total length (cm)

Length to tail (cm)

Tail length (cm)

Hind foot length (cm)

Chest circumferene (cm)

Rumpfat at 10 cm from the base of the tail (mm)

Rumpfat at 5 cm from the base of the tail (mm)

Rumpfat average (mm)

Peroneus muscle mass (g)

Antler spread (cm)

Number of points

Main antler beam (cm)

Distance to main beam (cm)

Spike length (cm)

**Table S2.** Estimation of heritability (*h*^2^) for body traits in white-tailed deer (*Odocoileus virginianus*). The genome relatedness matrices (GRM) were generated with varying minor allele frequency (MAF) and loci missing (LM) filters. Age and sex are included in the model but not shown. Standard errors for all metrics are included in parentheses and final filtered sample size is 399.

| GRM | MAF 1%  LM 70% | MAF 5%  LM 70% | MAF 10%  LM 70% | MAF 1%  LM 80% | MAF 1%  LM 90% | | MAF 1%  LM 100% | | MAF 5%  LM 80% | |
| --- | --- | --- | --- | --- | --- | --- | --- | --- | --- | --- |
| **Male & female GRM** |  |  |  |  |  | |  | |  | |
| Dressed body mass (kg) | 0.53 (0.24) | 0.45 (0.23) | 0.30 (0.14) | 0.52 (0.24) | | 0.54 (0.24) | | 0.28 (0.11) | | 0.45 (0.23) |
| Total length (cm) | 0.00 (0.22) | 0.00 (0.22) | 0.00 (0.12) | 0.00 (0.22) | | 0.00 (0.22) | | 0.07 (0.11) | | 0.00 (0.22) |
| Hind food length (cm) | 0.38 (0.26) | 0.37 (0.14) | 0.00 (0.14) | 0.37 (0.26) | | 0.35 (0.25) | | 0.07 (0.11) | | 0.36 (0.25) |
| Chest circumference (cm) | 0.26 (0.23) | 0.20 (0.14) | 0.18 (0.14) | 0.26 (0.23) | | 0.24 (0.23) | | 0.09 (0.11) | | 0.20 (0.22) |
| Peroneus muscle mass (g) | 0.57 (0.25) | 0.48 (0.14) | 0.39 (0.14) | 0.57 (0.25) | | 0.60 (0.25) | | 0.23 (0.11) | | 0.48 (0.24) |

**Table S3** Estimation of heritability (*h*^2^) for body traits in white-tailed deer (*Odocoileus virginianus*). The genomic-relatedness matrix (GRM) was generated using a minor allele frequency of 1% and required loci completion of 70%. Provide are *h*^2^ estimates with and without age and sex (denoted null) included in the model. Standard errors are included in parentheses.

| GRM | *Age + sex* | | *Null* | |
| --- | --- | --- | --- | --- |
|  | *N* | *h*^2^ | *N* | *h*^2^ |
| **Male & female GRM** |  |  |  |  |
| Dressed body mass (kg) | 399 | 0.53 (0.24) | 433 | 0.38 (0.22) |
| Total length (cm) | 399 | 0.00 (0.22) | 433 | 0.03 (0.20) |
| Hind food length (cm) | 399 | 0.38 (0.26) | 433 | 0.12 (0.20) |
| Chest circumference (cm) | 399 | 0.26 (0.23) | 433 | 0.14 (0.21) |
| Peroneus muscle mass (g) | 399 | 0.57 (0.25) | 433 | 0.35 (0.22) |

**Table S4.** Estimation of heritability (*h*^2^) for body traits in white-tailed deer (*Odocoileus virginianus*). The genomic-relatedness matrix (GRM) was generated using a minor allele frequency of 1% and required loci completion of 70%. Provide are *h*^2^ estimates with and without year of harvest included in the model. Standard errors are included in parentheses.

| GRM | *Age + sex + year* | | *Age + sex* | |
| --- | --- | --- | --- | --- |
|  | *N* | *h*^2^ | *N* | *h*^2^ |
| **Male & female GRM** |  |  |  |  |
| Dressed body mass (kg) | 399 | 0.49 (0.24) | 399 | 0.53 (0.24) |
| Total length (cm) | 399 | 0.00 (0.22) | 399 | 0.00 (0.24) |
| Hind food length (cm) | 399 | 0.39 (0.26) | 399 | 0.38 (0.26) |
| Chest circumference (cm) | 399 | 0.24 (0.23) | 399 | 0.26 (0.23) |
| Peroneos muscle mass (g) | 399 | 0.56 (0.25) | 399 | 0.57 (0.25) |

**Table S5.** Estimation of heritability (*h*^2^) for antler traits in white-tailed deer (*Odocoileus virginianus*). The genomic-relatedness matrix (GRM) was generated using a minor allele frequency of 1% and required loci completion of 70%. Provide are *h*^2^ estimates with and without year of harvest included in the model. Standard errors are included in parentheses.

| GRM | *Age + year* | | *Age* | |
| --- | --- | --- | --- | --- |
|  | *N* | *h*^2^ | *N* | *h*^2^ |
| **Male & female GRM** |  |  |  |  |
| Antler spread (cm) | 171 | 0.00 (0.54) | 171 | 0.22 (0.54) |
| No. of points | 182 | 0.52 (0.51) | 182 | 0.61 (0.51) |
| Main antler beam (cm) | 175 | 0.23 (0.50) | 175 | 0.21 (0.48) |

**Table S6.** Estimation of heritability (*h*^2^) for antler traits in white-tailed deer (*Odocoileus virginianus*). The genomic-relatedness matrix (GRM) was generated using a minor allele frequency of 1% and required loci completion of 70%. Mean trait values and effect sizes of age and SRP54 locus are provided; standard error for all metrics are included in parentheses. *N* refers to the number individuals retained after filtering for phenotype and genotype completeness.

| GRM | *N* | *h*^2^ | Mean | Age | SRP54 | |
| --- | --- | --- | --- | --- | --- | --- |
| **Male only GRM** | | | | | | |
| Antler spread (cm) | 71 | 0.99 (1.5) | 17 (2.0) | 3.3 (0.38) | | 1.6 (1.5) |
| No. of points | 74 | 0.99 (1.4) | 2.6 (0.44) | 0.64 (0.09) | | 0.39 (0.34) |
| Main antler beam (cm) | 71 | 0.99 (1.5) | 1.5 (0.11) | 0.18 (0.02) | | 0.00 (0.08) |
